# ANTIMICROBIAL SUSCEPTIBILITY PROFILE OF *Edwardsiella tarda* FROM FARMED FISH IN ILORIN METROPOLIS, KWARA STATE NIGERIA

**DOI:** 10.1101/2024.11.16.623970

**Authors:** F.O. Olabisi, I. Adeshina

## Abstract

This study determined the susceptibility profile and multiple antibiotic resistance (MAR) index of *Edwardsiella tarda* obtained from farmed fish in Ilorin Metropolis, Kwara State, Nigeria. 30 isolates were obtained from 30 catfish samples from the major fish market in Ilorin Metropolis. It was observed that the Isolates showed resistance to tetracycline (100%), ampicillin (100%), cefotaxime (100%) and ciprofloxacin (97%). *Edwardsiella tarda* isolates were entirely susceptible to Gentamycin (100%) while also showing varying levels of susceptibility to Enfloxacin (77%), Norfloxacin (67%), Sulphamethoxazole (50%), Composulphonamides (23%) and Chloramphenicol (10%). The isolates showed an intermediate range of susceptibility to Chloramphenicol (90%), Composulphonamides (77%), Sulphamethoxazole (50%) and Ciprofloxacin (3%). Analysis of the MAR index of isolates revealed that 97% of the isolates had a MAR index of 0.4, with only 3% of the isolates showing a multiple antibiotic resistance index of 0.3, which is an indication that farmed fishes in the Ilorin metropolis had a high risk of exposure to antibiotics.

## 1.0 INTRODUCTION

Antibiotic abuse and uncontrolled use in the fish farming industry have significantly increased the amount of resistant microorganisms in the world including Nigeria. These antimicrobial practices have been widely used in fish farming industry exclusively for therapeutic or preventative purposes (Manyi-Loh *et al*., 2018). However, these antibiotics are typically used to boost and speed up fish growth and for effective feed efficiency in the fish (Muhammad *et al*., 2020).

Antibiotics are widely used in Nigerian aquaculture and livestock husbandry. All antibiotic resistance in microbes has a genetic basis (Abu & Wondikom, 2018) since it is a defence mechanism that enables bacteria to survive in an unfavourable environment (Gang *et al*., 2015).

Antibiotic resistance represents one of the most significant current threats to public health and is predicted to overtake cancer as a cause of death by 2050. Nevertheless, little or no attention has been paid to the potential use of antibiotics in aquaculture industries. In addition to the transfer of antibiotic-resistant microorganisms and their genes through consuming contaminated fish, there is a significant risk of environmental contamination due to using medicated feeds.

Antibiotic resistance has been reported in catfish, shrimp, eels, and aquaculture environments. According to some accounts, resistance develops shortly after using antibacterial medications to treat infections, reducing their effectiveness in managing bacterial fish diseases. Antimicrobial usage must, therefore, be carefully regulated and monitored in aquaculture and our environment (Ma *et al*., 2021) to evaluate their impact and subsequent risk to the ecosystem.

*Edwardsiella tarda* is the causative agent of Edwardsiellosis in fish, characterised by systemic hemorrhagic septicemia, internal abscesses and skin lesions (Ewing *et al*., 1965). The bacterium is opportunistic, and disease outbreaks occur when poor environmental conditions prevail, like overcrowding, poor water quality conditions, high organic content and high temperature (Mohanty and Sahoo, 2007).

This study investigated the occurrence of antimicrobial resistance of *Edwardsiella tarda* from farmed fish within Ilorin metropolis in Kwara State to identify high-risk sources of contamination and address the issue of emerging antibiotic resistance of *Edwardsiella tarda* bacteria from fish farming.

## 2.0 MATERIALS AND METHODS

### 2.1 Description of Study Area

The study area is within the Ilorin metropolis, Kwara State, and is characterised by various economic activities such as fish farming, trading, and fishing. It is also an important industrial and commercial area with many local manufacturing and trade companies.

### 2.2 Collection of Fish Samples

30 fish samples were collected from the major fish market in Ilorin Metropolis, Kwara State. Fish samples were aseptically dissected using a dissecting kit.

### 2.3 Isolation and Identification of *Edwardsiella tarda*

➢ Swabs were taken aseptically from the intestine of the dissected fish samples and were streaked on Petri dish-containing Tryptone soy agar (LAB M Ltd., United Kingdom) that was prepared according to manufacturers’ instructions.
➢ The plates were incubated at 37 °C for 24 hours and were observed for bacterial growth and distinct colonies were further sub-cultured on freshly prepared tryptone soy agar to obtain a pure culture of the isolates.
➢ *Edwardsiella tarda* isolates were then identified using gram staining technique, morphology, motility, and biochemical reaction tests such as oxidase test, catalase test, lactose fermentation and glucose fermentation according to (Bergey’s Manual of Determinative Bacteriology, 7th Edition).
➢ Measurement of the zone of inhibition (zi), The zone of inhibition was measured using meter rule in millimeters(mm).

### 2.4 Antibiotic susceptibility testing

Antibiotic Susceptibility testing was carried out with a commercially available antimicrobial susceptibility test single disc from (Oxoid Ltd, Basingstoke, RG24 8PW, United Kingdom) Pure culture of *Edwardsiella tarda* was picked from Tryptone soy broth using a sterile wire loop and transferred to tubes each containing 5 ml of sterile physiological saline.

The suspension was vortexed and adjusted to match 0.5 McFarland turbidity standards. Sterile swab sticks were then dipped, rotated and pressed firmly on the tube walls above the culture to remove excess inoculums from the swabs.

This was then evenly swabbed on the dried surface of Mueller-Hinton agar (Oxoid Ltd., England) plates ensuring even distribution of the bacterium.

10 different antimicrobial susceptibility single discs were placed on the plates using sterile forceps and allowed to undergo incubation at a temperature of 37 °C for 18 to 24 hours, after which the diameter of the zone of inhibition was measured with a white transparent meter rule in millimetre.

The antimicrobial susceptibility test was carried out using the agar disk diffusion method as described by Bauer *et al*. (1966)

The methods described above and the interpretations of results were done using the Clinical Laboratory Standard Institute (CLSI. 2020) manual.

The ten antibiotics that were used have the following concentrations: Enfloxacin (ENR) (5μg), Ampicillin (AMP) (10μg), Norfloxacin (NOR) (10μg), Gentamycin (CN) (10μg), Cefotaxime (CTX) (5μg), Tetracycline (TE) (10μg), Composulphonamides (S3) (30μg), Sulphamethoxazole (SXT) (5μg), Chloramphenicol (C) (30μg), Ciprofloxacin (CIP) (5μg).

### 2.5 Determination of Antibiotics Resistance Index

In vitro sensitivity test was done on random selected isolates of *Edwardsiella tarda* using disc diffusion method according the method carried out by Bauer *et al*. (1966) using different antimicrobial agents: Enfloxacin (ENR) (5μg), Ampicillin (AMP) (10μg), Norfloxacin (NOR) (10μg), Gentamycin (CN) (10μg), Cefotaxime (CTX) (30μg), Tetracycline (TE) (10μg), Composulphonamides (S3) (30μg), Sulphamethoxazole (SXT) (25μg), Chloramphenicol (C) (30μg), Ciprofloxacin (CIP) (5μg).

The diameter of the inhibition zone was measured and interpreted according to clinical and laboratory standards institute (CLSI, 2020) manual. Isolates that showed resistance to more than two different antibiotic groups were multiple drug-resistant (MDR) isolates.

### 2.6 Determination of Multiple Antibiotics Resistance (MAR) Index

Multiple antibiotics resistance (MAR) index is an analytical measure or technique of analysis introduced in 1983 by Krumperman which helps to categorize or group different sources of high and low-risk contamination of bacterial isolates based on their frequency of antibiotic resistance (Osundiya *et al*., 2013).

Multiple Antibiotic Resistance (MAR) index for each strain was identified according to Singh *et al*., (2010): MAR index = No. of resistance (Isolates classified as intermediate were considered sensitive for MAR index) / Total No. of tested antibiotics The multiple antibiotics index is a measure used to identify the risk of contamination that poses a potential hazard to humans. However, multiple antibiotics resistance value greater than 0.2 is a signal that the bacterial isolates were from high-risk sources while bacterial isolates with multiple antibiotics resistance index less than 0.2 is an indicator of low-risk contamination. (Nyandjou *et al*., 2019).

### 2.7 Interpretation of the data obtained with Clinical Laboratory Standard Institute, CLSI (2020) Manual

The Clinical Laboratory Standard Institute Manual 2020 was used for the interpretation of the diameter (mm) of the zone of inhibition of the antimicrobials carried out during the experimental work.

**Table 2.8:** Clinical Laboratory Standard Institute, CLSI (2020) Manual.

| **Zone of Inhibition (mm)** | **Susceptible (S)** | **Intermediate (I)** | **Resistant (R)** |
| --- | --- | --- | --- |
| Enfloxacin (ENR) | $\geq 18$ | 15 – 17 | $\leq 14$ |
| Gentamycin (CN) | $\geq 15$ | 13 – 14 | $\leq 12$ |
| Chloramphenicol (C) | $\geq 18$ | 13 – 17 | $\leq 12$ |
| Ciprofloxacin (CIP) | $\geq 26$ | 22 – 25 | $\leq 21$ |
| Tetracycline (TE) | $\geq 15$ | 12 – 14 | $\leq 11$ |
| Ampicillin (AMP) | $\geq 17$ | 14 – 16 | $\leq 13$ |
| Norfloxacin (NOR) | $\geq 17$ | 13 – 16 | $\leq 12$ |
| Cefotaxime (CTX) | $\geq 26$ | 23 – 25 | $\leq 22$ |
| Composulphonamides(S3) | $\geq 17$ | 13 – 16 | $\leq 12$ |
| Sulphamethoxazole (SXT) | $\geq 16$ | 11 – 15 | $\leq 10$ |

### 2.8 Statistical Analysis

The data (diameter of inhibition zone) was measured and interpreted according to clinical and laboratory standards institute (CLSI, 2020) manual. The data obtained was analyzed using descriptive statistics with the aid of SPSS version 25.

## 3.0 RESULTS

### 3.1 Susceptibility profile of *Edwardsiella tarda* Isolates (n = 30) from farmed fishes to 10 antibiotics

The data (diameter of inhibition zone) was measured and interpreted according to clinical and laboratory standards institute (CLSI, 2020) manual. The table below shows the susceptibility percentage of *Edwardsiella tarda* isolates tested with ten antibiotics, it can be inferred that all 30 isolates showed a 100% resistance to tetracycline, ampicillin and cefotaxime antibiotics while showing a 97% resistance to ciprofloxacin.

Whereas the isolates showed a 0% resistance to Enfloxacin, Gentamycin, Norfloxacin, Chloramphenicol, Composulphonamides, and Sulphamethoxazole. Gentamycin was 100% susceptible to all the isolates while Enfloxacin, Norfloxacin, Sulphamethoxazole and Composulphonamides showed 77%, 67%, 50% and 23% susceptibility respectively. Chloramphenicol Composulphonamides, Sulphamethoxazole, Enfloxacin, Norfloxacin, and Ciprofloxacin also showed an intermediate range of 90%, 77%, 50%, 33%, 23% and 3% respectively.

**TABLE 3.2:**
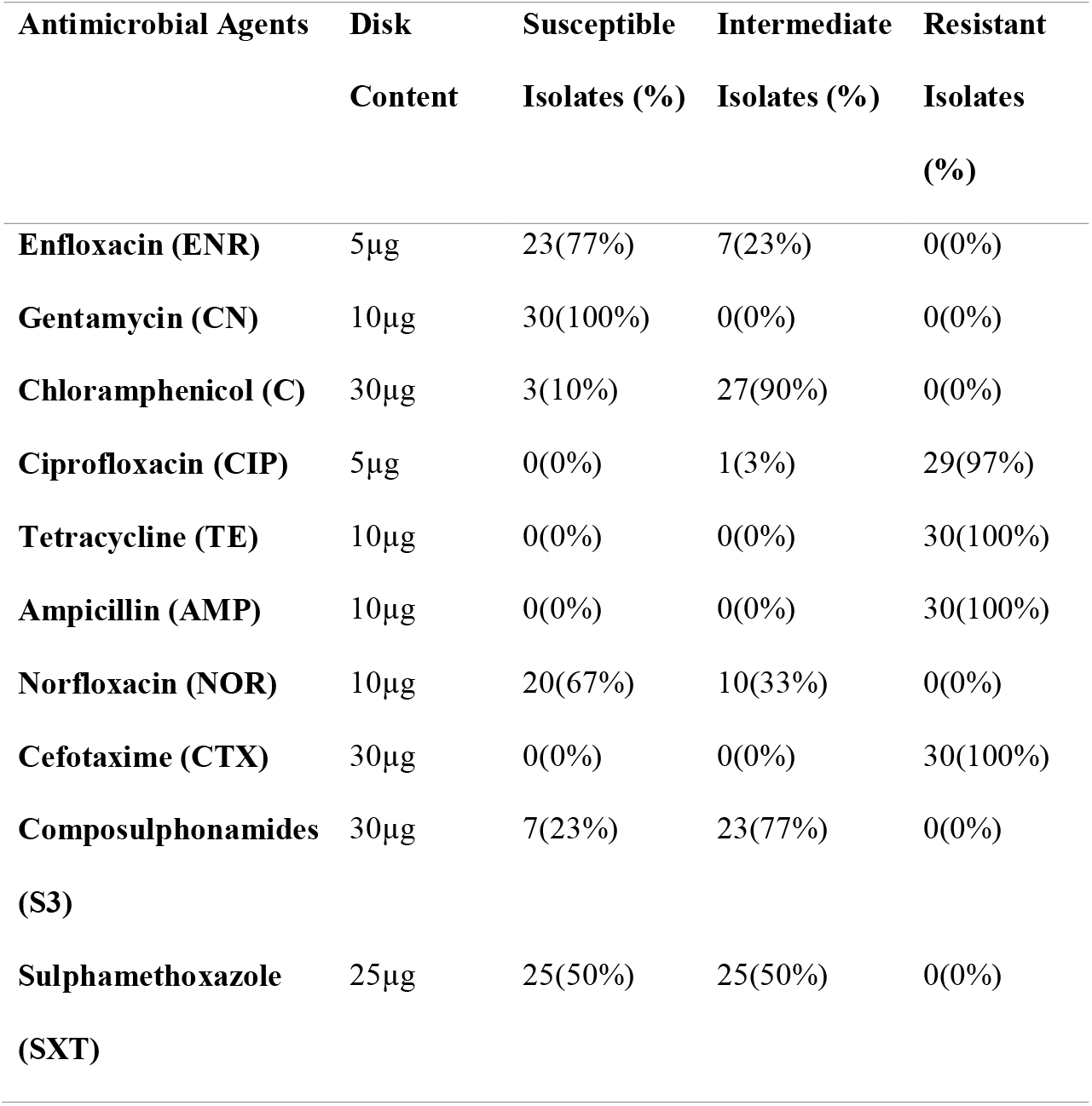
Susceptibility profile of *Edwardsiella tarda* Isolates (n = 30) from farmed fishes to 10 antibiotics.

| **Antimicrobial Agents** | **Disk** | **Susceptible** | **Intermediate** | **Resistant** |
| --- | --- | --- | --- | --- |
|  | **Content** | **Isolates (%)** | **Isolates (%)** | **Isolates (%)** |
| **Enfloxacin (ENR)** | 5µg | 23(77%) | 7(23%) | 0(0%) |
| **Gentamycin (CN)** | 10µg | 30(100%) | 0(0%) | 0(0%) |
| **Chloramphenicol (C)** | 30µg | 3(10%) | 27(90%) | 0(0%) |
| **Ciprofloxacin (CIP)** | 5µg | 0(0%) | 1(3%) | 29(97%) |
| **Tetracycline (TE)** | 10µg | 0(0%) | 0(0%) | 30(100%) |
| **Ampicillin (AMP)** | 10µg | 0(0%) | 0(0%) | 30(100%) |
| **Norfloxacin (NOR)** | 10µg | 20(67%) | 10(33%) | 0(0%) |
| **Cefotaxime (CTX)** | 30µg | 0(0%) | 0(0%) | 30(100%) |
| **Composulphonamides (S3)** | 30µg | 7(23%) | 23(77%) | 0(0%) |
| **Sulphamethoxazole (SXT)** | 25µg | 25(50%) | 25(50%) | 0(0%) |

### 3.3 Multiple Antibiotic Resistance (MAR) Index of *Edwardsiella tarda* strains (n = 30) isolates in this study

The table below shows the number of resistant antibacterial from all *Edwardsiella tarda* isolates (n = 30) and the multiple antibiotic resistance index of all the isolates. Multiple Antibiotic Resistance index is used to determine the high risk of contamination with the use of antibiotics that pose a high risk to humans and also the prevalence of antibiotics in the treatment of *Edwardsiella tarda* bacteria.

Multiple Antibiotic Resistance (MAR) index for each isolate was identified according to Singh *et al*., (2010): MAR index = Number of resistant antibiotics in a single isolate (Number of antibiotics to which the bacterial isolates showed resistance) / Total Number of tested antibiotics. A Multiple Antibiotic Resistance index value of equal to or less than 0.2 was defined as those antibiotics that were seldom or never used for the animal in terms of treatment, whereas a MAR index value higher than 0.2 is considered that the fish has received high-risk exposure to those antibiotics.

Therefore, it can be deduced from the table below that the fishes had received a high risk of exposure to antibiotics commonly used in aquaculture practices as all the isolates showed a MAR index ranging from 0.3 to 0.4. All isolates showed a MAR index of 0.4 except for F27 which showed a MAR index of 0.3 with a total of three antibiotics showing resistance to the isolate. The average MAR index for all the tested isolates was 0.397.

**Table 3.4:** Multiple Antibiotic Resistance (MAR) Index of *Edwardsiella tarda* strains (n = 30) isolates in this study.

| **Isolates** | **No resistant**<br>**antibacterial** | **MAR index** |
| --- | --- | --- |
| **F1** | 4 | 0.4 |
| **F2** | 4 | 0.4 |
| **F3** | 4 | 0.4 |
| **F4** | 4 | 0.4 |
| **F5** | 4 | 0.4 |
| **F6** | 4 | 0.4 |
| **F7** | 4 | 0.4 |
| **F8** | 4 | 0.4 |
| **F9** | 4 | 0.4 |
| **F10** | 4 | 0.4 |
| **F11** | 4 | 0.4 |
| **F12** | 4 | 0.4 |
| **F13** | 4 | 0.4 |
| **F14** | 4 | 0.4 |
| **F15** | 4 | 0.4 |
| **F16** | 4 | 0.4 |
| **F17** | 4 | 0.4 |
| **F18** | 4 | 0.4 |
| **F19** | 4 | 0.4 |
| **F20** | 4 | 0.4 |
| **F21** | 4 | 0.4 |
| **F22** | 4 | 0.4 |
| **F23** | 4 | 0.4 |
| **F24** | 4 | 0.4 |
| **F25** | 4 | 0.4 |
| **F26** | 4 | 0.4 |
| **F27** | 3 | 0.3 |
| **F28** | 4 | 0.4 |
| **F29** | 4 | 0.4 |
| **F30** | 4 | 0.4 |
| **Average** |  | 0.397 |

**Plate 3.5:**
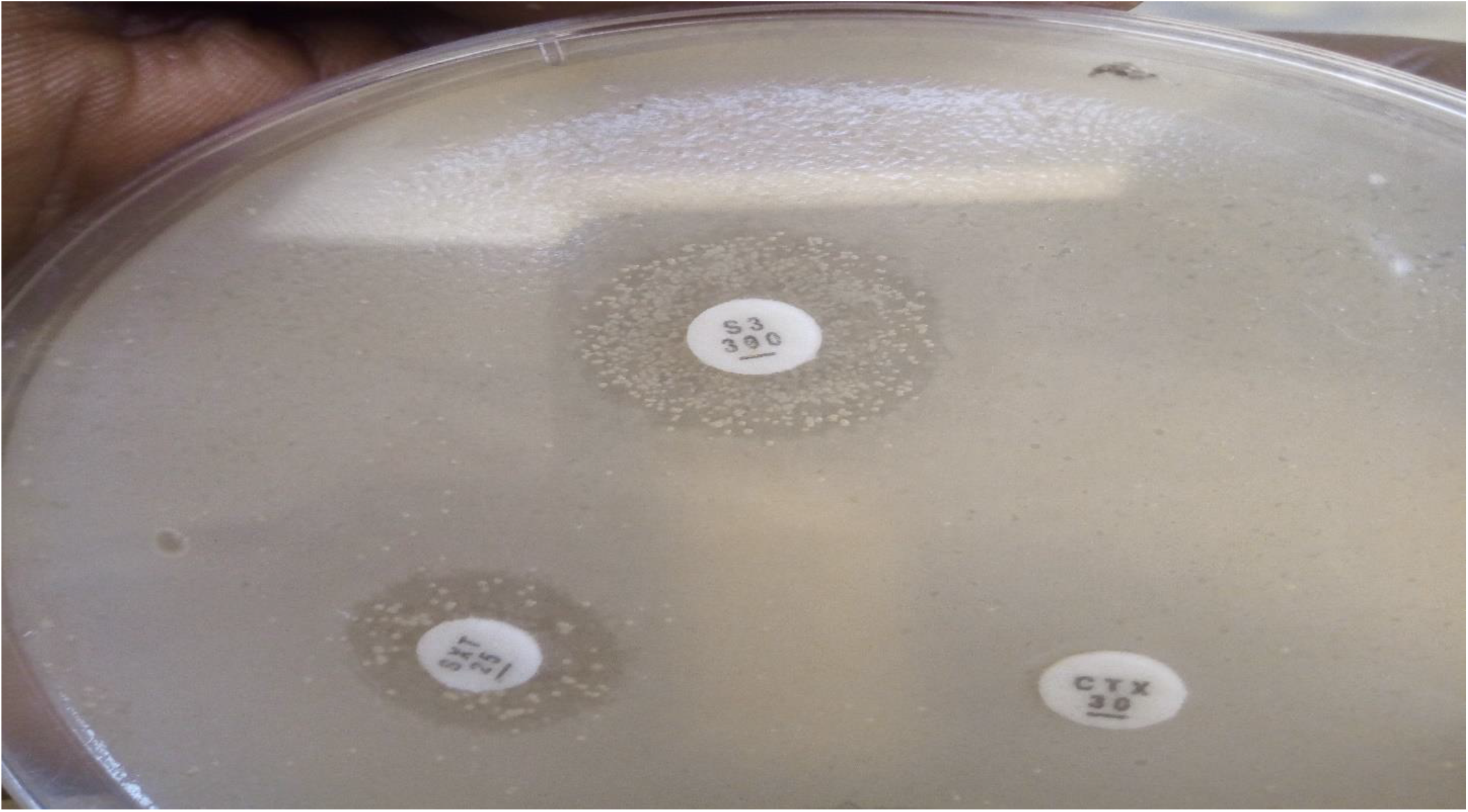
Antimicrobial susceptibility disk containing Sulphamethoxazole (SXT), Composulphonamides (S3) and Cefotaxime (CTX) Antibiotics containing a disk content of 25μg, 30μg and 30μg respectively.

**Plate 3.6:**
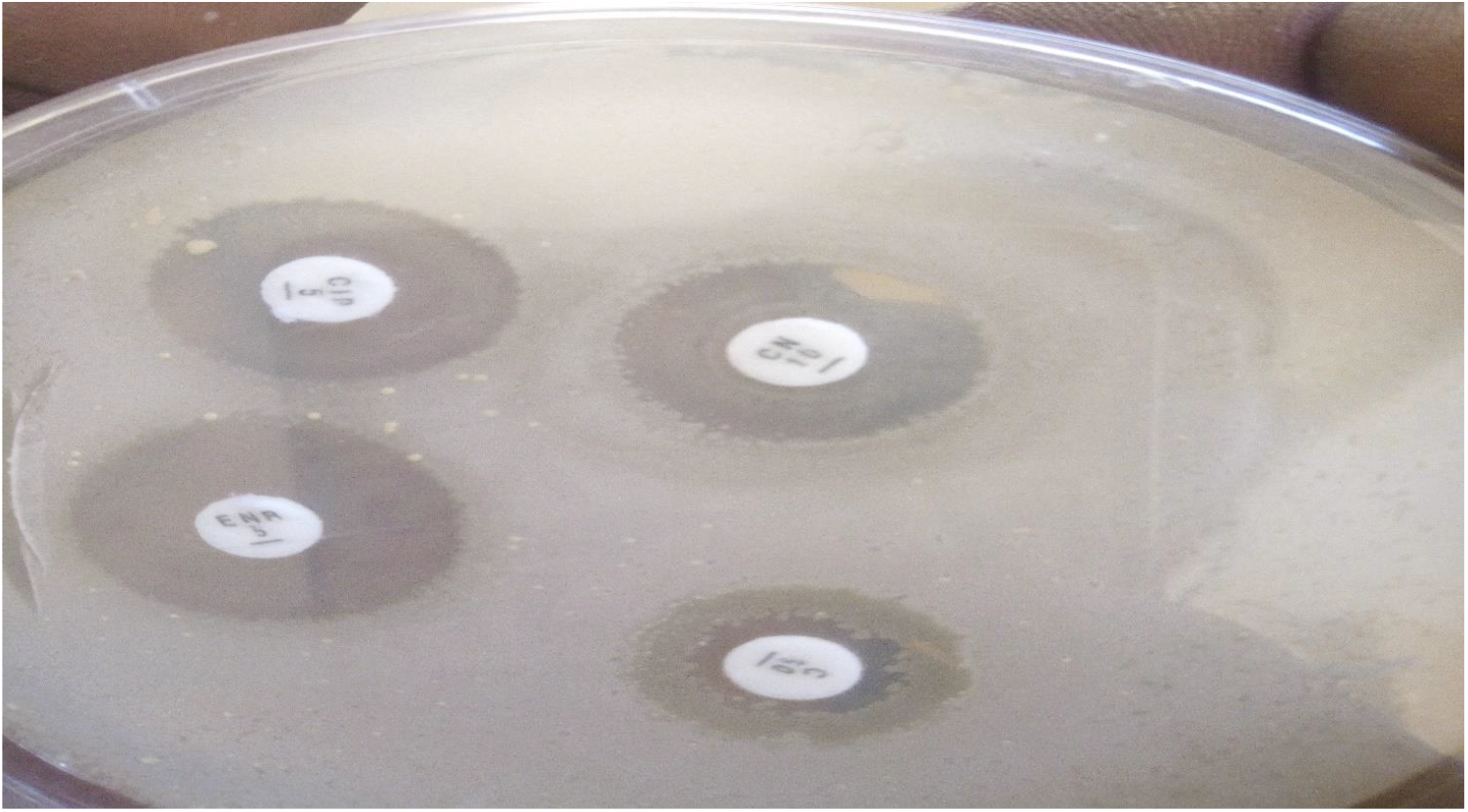
Antimicrobial susceptibility disk containing Enfloxacin (ENR), Gentamycin (CN), Chloramphenicol (C) and Ciprofloxacin (CIP) Antibiotics containing a disk content of 5μg, 10μg, 30μg and 5μg respectively.

**Plate 3.7:**
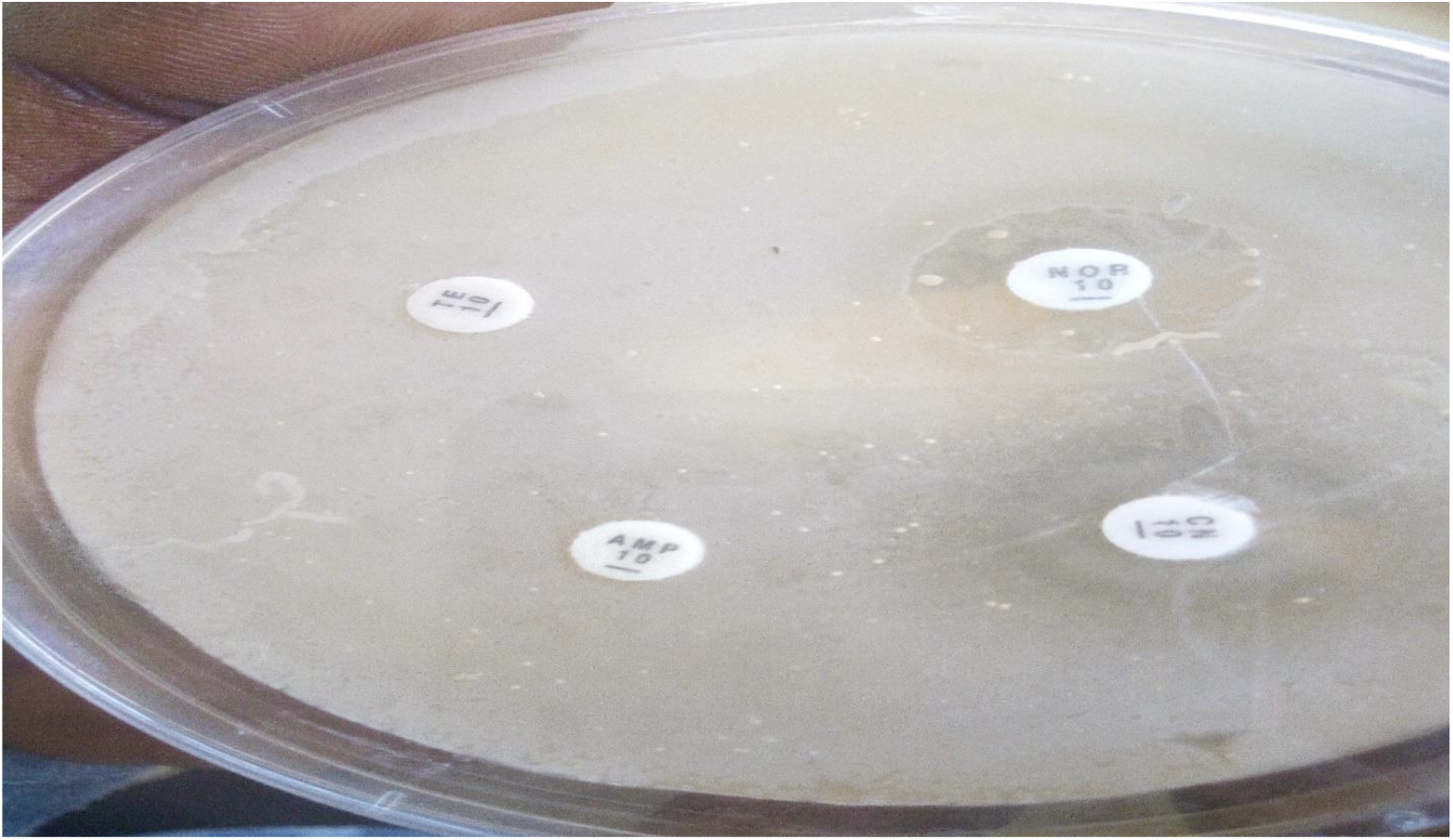
Antimicrobial susceptibility disk containing Tetracycline (TE), Ampillicin (AMP), and Norfloxacin (NOR) Antibiotics containing a disk content of 10μg, 10μg, and 10μg respectively

### 3.8 Antimicrobial Susceptibility Profile of *Edwardsiella tarda* Isolates (n = 30) isolated in this study also show their multiple antibiotic resistance index (MAR)

The table below shows the susceptibility profile of *Edwardsiella tarda* isolates to ten commonly used antibiotics in aquaculture, the multiple antibiotics resistance index (MAR) was also determined from the number of antibiotics that were resistant to the isolate(s)/ total number of antibiotics tested.

MAR index greater than 0.2 is considered that the fish had received high-risk exposure to those antibiotics. Therefore, 97% of *Edwardsiella tarda* isolates showed an MAR index of 0.4 while only 3% of *Edwardsiella tarda* isolates showed a MAR of 0.3 indicating that the fish were highly exposed to the antibiotics that were tested on the isolates.

The mean value of the MAR index showed a value of 0.397, *Edwardsiella tarda* isolates showed complete resistance to tetracycline, ampicillin and cefotaxime while showing a 90% resistance to chloramphenicol.

*Edwardsiella tarda* was susceptible to Gentamycin, Enfloxacin and Norfloxacin, indicating that Gentamycin, Enfloxacin and Norfloxacin are effective in controlling *Edwardsiella tarda* infections.

**Table 3.9:**
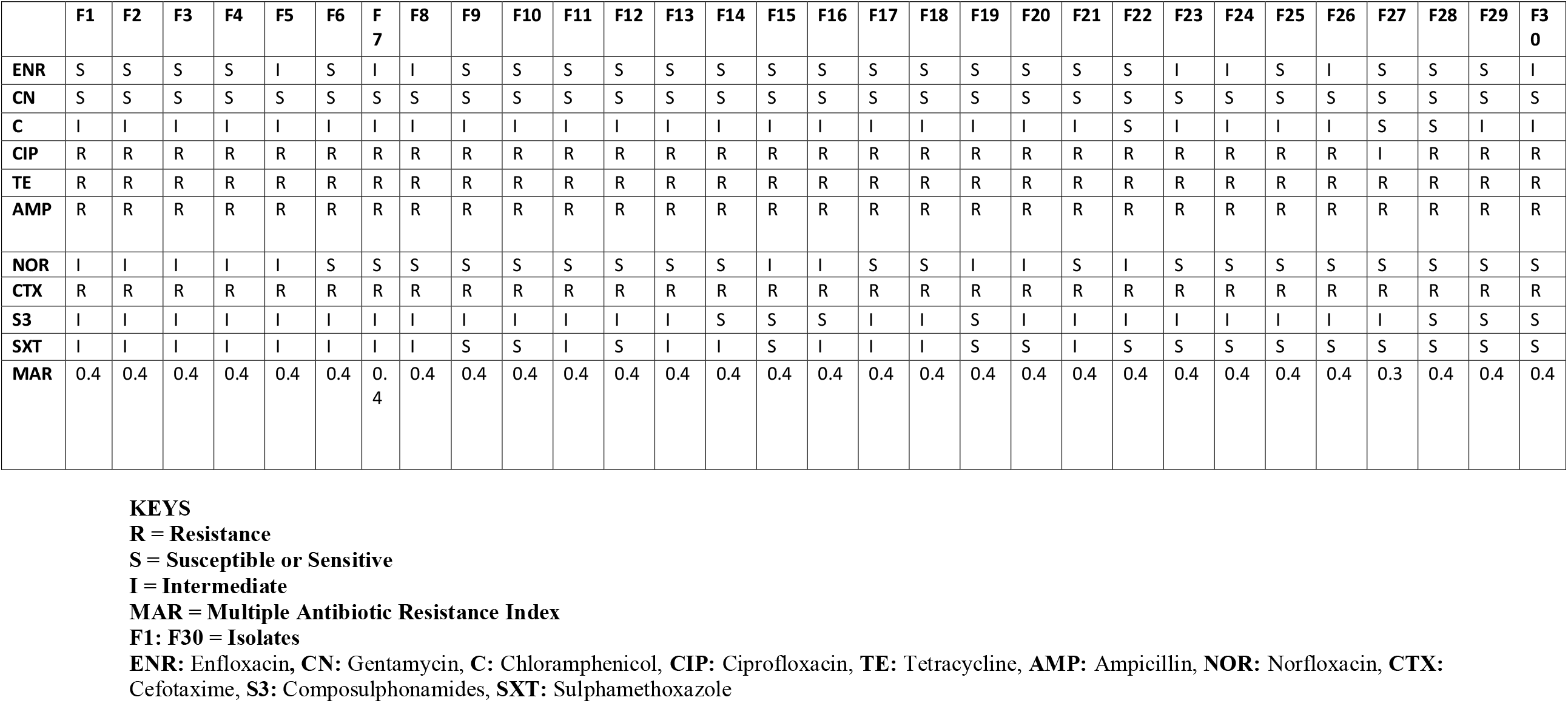
Antimicrobial Susceptibility Profile of *Edwardsiella tarda* Isolates (n = 30) isolated in this study also showing their multiple antibiotic resistance index (MAR)

## 4.0 DISCUSSION AND CONCLUSION

### 4.1 Discussion

Antimicrobial agents can be used as a tool to maintain the health and disease prevention of cultured animals (Bischoff *et al*., 2005). However, overuse of antimicrobial drugs has the potential to increase the occurrence of antibiotic resistance in harmful bacteria, lowering their susceptibility to antibiotics.

Therefore, this study was carried out to reveal the antimicrobial resistance pattern and antimicrobial resistance prevalence of *Edwardsiella tarda*, an important bacterial disease in freshwater fish.

*Edwardsiella tarda* is considered as one of the most important bacterial microorganisms that cause severe economic losses due to morbidity and mortality among various populations and age groups of fish in many countries. It is considered a dangerous septicemic pathogen with high economic losses and highly encountered causes of diseases in stressed warm water aquaculture (Jun & Yin, 2006; Ibrahem *et al*., 2011).

Antimicrobial sensitivity test in this study revealed that isolated *Edwardsiella tarda* was 100% resistant to Tetracycline, Ampilicin and Cefotaxime while 29 of the *Edwardsiella tarda* isolates showed 100 % resistance to Ciprofloxacin with a total resistance of 97%, Cefotaxime showed a resistance to *Edwardsiella tarda* isolates which was also agreed to (Nagy *et al*., 2018) from their study justifying the case that cefotaxime is also a resistant antibiotic.

This result indicates multiple antibiotic resistance and a total resistance to commonly used antibiotics in aquaculture such as Tetracycline, Ampilicin and Ciprofloxacin which are the frequently administered antibiotics used for treating infections in humans and also in aquaculture.

Wimalasena *et al*., (2018) reported that resistant isolates were at high frequency for the β-lactams antibiotics which are ampicillin and cefotaxime but at low frequency for the aminoglycosides (tetracycline) while Abd El Tawab *et al*. (2020) recorded a 75% resistance to ampicillin.

Ogbonne *et al*. (2018) reported that *Edwardsiella tarda* showed a 100% resistance to Amoxycillin, which is under the same group of β-lactams antibiotics.

*Edwardsiella tarda* isolates were also reported to be 100% resistant to chloramphenicol by Ogbonne et al., (2018), which was in contrast with this study, where chloramphenicol showed no resistance to *Edwardsiella tarda* but a 90% intermediate range to *Edwardsiella tarda* which is similar to the work of (Nagy *et al*., 2018).

Abd El Tawab *et al*. (2020) reported that Ciprofloxacin was 100% susceptible to *Edwardsiella tarda*, which was in agreement with (Ogbonne *et al*., 2018). Whereas, the study here showed that Ciprofloxacin was 96.67% resistant to *Edwardsiella tarda*, with an indication that E.tarda is highly resistant to Ciprofloxacin.

Tetracycline was found to be 100% resistant to *Edwardsiella tarda* in this study, while Abd El Tawab *et al*. (2020) reported 25% resistance of tetracycline to *Edwardsiella tarda* isolates in Egypt.

Sulphamethoxazole showed a 50% intermediate-range and 50% sensitivity range to *Edwardsiella tarda*, which was different from the study of (Nagy et al., 2018), where the antibiotic was resistant to the isolates. Abd El Tawab *et al*., (2020) reported that Sulphamethoxazole showed a 100% susceptible range to *Edwardsiella tarda* which was different from the result in this study where Sulphamethoxazole showed a 50% intermediate sensitivity range and 50% sensitivity range to *Edwardsiella tarda* indicating that the bacteria is developing resistance to the antibiotic over time due frequent and uncontrolled usage.

The study carried out shows that Gentamycin, Enfloxacin and Norfloxacin were susceptible to *Edwardsiella tarda* isolates which were in close agreement with the result of Ogbonne *et al*., (2018) and Abd El Tawab *et al*., (2020). This therefore implies that Gentamycin, Enfloxacin and Norfloxacin are effective in controlling *Edwardsiella tarda* infections.

However, our results showed resistance to more than one class of antibiotic, and this is in agreement with the work of (Lee et al. 2011), whose study reported multiple drug resistance by *Edwardsiella tarda* in freshwater fish, indicating that the farmed fishes within the Ilorin metropolis have high exposure to antibiotics.

### 4.2 Conclusion

The result of the present study shows the isolation of *Edwardsiella tarda* from commonly sold farmed fish in Ilorin Metropolis; the study also shows that *Edwardsiella tarda* is resistant to some commonly used antibiotics in aquaculture practices.

The study shows the prevalence of antimicrobial resistance in *Edwardsiella tarda* from farmed fishes in Ilorin Metropolis, indicating that farmed fishes in Ilorin Metropolis had a high risk of exposure to antibiotics.

